# ACE2 peptide fragment interacts with several sites on the SARS-CoV-2 spike protein S1

**DOI:** 10.1101/2020.12.29.424682

**Authors:** Aleksei Kuznetsov, Piret Arukuusk, Heleri Härk, Erkki Juronen, Ülo Langel, Mart Ustav, Jaak Järv

## Abstract

The influence of the peptide QAKTFLDKFNHEAEDLFYQ on the kinetics of the SARS-CoV-2 spike protein S1 binding to angiotensin-converting enzyme 2(ACE2) was studied to model the interaction of the virus with its host cell. This peptide corresponds to the sequence 24-42 of the ACE2 α1 domain, which is the binding site for the S1 protein. The on-rate and off-rate of S1-ACE2 complex formation were measured in the presence of various peptide concentrations using Bio-Layer Interferometry (BLI). The formation of the S1-ACE2 complex was inhibited when the S1 protein was preincubated with the peptide, however, no significant inhibitory effect was observed in the absence of preincubation. Dissociation kinetics revealed that the peptide remained bound to the S1-ACE2 complex and stabilized this complex. Computational mapping of the S1 protein surface for peptide binding revealed two additional sites, located at some distance from the receptor binding domain (RBD) of S1. These additional binding sites affect the interaction between the peptide, the S1 protein, and ACE2.

## Introduction

Binding of SARS-CoV-2 particles to the angiotensin-converting enzyme 2 (ACE2) on host cells leads to the fusion of the virus and host cell membranes and initiates the entrance of the viral RNA into the cells [1-2]. It has been suggested that blocking this binding process will inhibit the virus entry process and thus may have a therapeutic antiviral effect [3]. The most straightforward way of designing inhibitors is to use peptides to mimic the interaction interface between the spike protein and the ACE2 molecule in complex [4-6]. The spike protein is composed of S1and S2 domains, and it is S1 that contains the receptor-binding domain (RBD) that binds to ACE2 [7-8]. As the molecular structure of the S1-ACE2 complex is known [7-9], and the atomic coordinates and experimental data (code 6LZG) have been deposited in the PDB database (*www.wwPDB.org*), inhibitory peptides can be designed based on the structure of the complex. However, as ACE2 is a physiologically important enzyme, its inhibition by antiviral prophylaxis with peptides derived from the spike protein is not a promising approach. Therefore, we designed peptides that are derived from the ACE2 structure and interact with the RBD of the spike protein S1 [5, 6].

Initially, the ACE2 binding site on the S1 protein was mapped computationally [5]. This analysis revealed that the peptide STIEEQAKTFLDKFNHEAEDLFYQSSL, derived from the α1 domain of the N-terminal part of ACE2, containing amino acids 19-45, can be truncated from both ends without significant loss of binding. Therefore, the shorter peptide QAKTFLDKFNHEAEDLFYQ (amino acids 24 - 42), which still interacts effectively with the S1 protein (docking energy E_dock_ = −11.7 kcal/mol [5]), was selected for experiments in this study. The molecular mass of this peptide is too low for a direct binding assay using Bio-Layer Interferometry technology (BLI) [10], and it is unclear how chemical modification or loading with a cargo molecule or linker group would influence the binding properties of the peptide. Therefore, we loaded ACE2 onto the biosensors and studied the influence of the peptide on the kinetics of the S1-ACE2 interaction.

## Materials and methods

### Peptide synthesis

The peptide QAKTFLDKFNHEAEDLFYQ was synthesized on an automated peptide synthesizer (Biotage Initiator+ Alstra, Sweden) using the fluorenylmethyloxycarbonyl (Fmoc) solid-phase peptide synthesis strategy and Rink-amide ChemMatrix resin (PCAS-BioMatrix, Québec, Canada) to obtain C-terminally amidated product. N,N’-diisopropylcarbodiimide (DIC) and Oxyma Pure in dimethylformamide (DMF) were used as coupling reagents, and N, N-diisopropylethylamine (DIEA) was used as the activator base. Cleavage of the product was performed with trifluoroacetic acid (TFA), 2.5% triisopropylsilane (TIPS), and 2.5% water for three hours at room temperature.

The peptide was purified by high-performance liquid chromatography (HPLC) on a C4 column (Phenomenex Jupiter C4, 5 μm, 300 Å, 250 × 10 mm, Agilent) using an acetonitrile/water gradient containing 0.1% TFA. The purity of the peptide was validated at 98% using a Waters Acquity Ultra-Performance Liquid Chromatography (UPLC) with an acetonitrile/water gradient (Supplement Fig S1). The accurate molecular weight of the peptide was determined to be 2342 Da using a matrix-assisted laser desorption/ionization time-of-flight (MALDI-TOF) mass spectrometer (Brucker Microflex LT/ SH, USA), with α-cyano-4-hydroxycinnamic acid as the matrix (Supplement Fig S2). The calculated molecular weight of the peptide was 2342.15 Da.

### Proteins

Human recombinant ACE2-His protein (Icosagen OÜ, Estonia, cat# P-302-100) and SARS-CoV-2 Spike protein S1 (Icosagen OÜ, Estonia, cat# P-305-100) were used in this study.

### Bio-Layer interferometry (BLI)

His-tagged ACE2 was immobilized onto Octet RED96e biosensors (ForteBio, CA, USA) and the binding of S1 protein was measured in the presence or absence of peptide QAKTFLDKFNHEAEDLFYQ. Experiments were performed at 25◻°C in 20 mM Tris-HCl pH 7.0 and 150 mM NaCl. Biosensors (HIS1K, lot 6110102) were loaded with His-tagged ACE2, before S1 protein or S1 protein and peptide were added to start the complex formation process. In one series of experiments, the peptide was preincubated with S1 for 15 min at 25◻°C before the binding assay. Complex dissociation was initiated by immersing the biosensors into fresh assay buffer (20 mM Tris-HCl pH 7.0 with 150 mM NaCl), without S1 protein and peptide. Data were analyzed using ForteBio Data Analysis software (version 11.1.1.39) [11]. Results are presented in the Supplement Table S1 and Table S2.

### Computational peptide docking

The peptide docking study was performed as described previously [5,6]. Briefly, the input file for modeling the S1-ACE2 complex was built from X-ray analysis data [7,9] deposited in the PDB database (*www.wwPDB.org*) with the code 6LZG. GROMACS package (version 4.6.1) was used for molecular dynamics simulations [12] and AutoDock Vina (version 1.1.2) was used for ligand docking [13]. The best scores were selected for peptide positioning. Protein protonation at pH 7 was processed using the GROMACS pdb2gmx tool, and the geometric, charge, and van der Waals constrained parameters were assigned using the GROMOS 53a6 force field parameter set [14]. The protein structure, neutralized by adding Na^+^ and Cl^−^ ions, was solvated in a 5 nm cubic box, filled with SPC water as solvent [15]. The system was allowed to reach equilibrium at constant pressure (1 atm) [16] and temperature (300 K), controlled by the modified Berendsen thermostat algorithm [17]. Equilibrated simulations were performed on the systems for ten nanoseconds. After MD relaxation, the protein structure was extracted from the system and used for docking procedures. The docking compatible structure formats of the protein were prepared by AutoDockTools (version 1.5.6) [18]. The fitting box with 0.3 Å of grid spacing was defined once and used for all docking calculations. The fitting area covered the whole protein space and the docking poses were obtained and listed following the docking energy values. The graphic software package VMD (version 1.9.4) was used to illustrate ligand docking poses on the protein surface [19].

## Results and discussion

### Kinetic measurements of the S1-ACE2 binding interaction

The influence of the peptide QAKTFLDKFNHEAEDLFYQ on the binding of the SARS-CoV-2 spike S1 protein with ACE2 was investigated by loading ACE2 onto biosensors, then dipping into a buffer containing S1 protein, or S1 protein and peptide. This experimental setup allowed characterization of the complex formation and dissociation reactions, described by the ascending and descending parts of the graphs, respectively, exemplified in Fig 1. Taking the ascending part of the plot, the complex formation reaction is characterized by the first-order rate constant k_on_ (s^−1^) and the second-order rate constant k_ass_ (M^−1^s^−1^). In the latter case, the concentration of the S1 protein in the assay buffer is taken into consideration [10]. The descending part of the plot allows calculation of the complex dissociation rate constant, denoted here as k_off_ (s^−1^). The equilibrium constant, K_D,_ for the complex dissociation can be calculated as the ratio of the k_off_ and k_ass_ values [10, 11]. In this study, the K_D_ values for the S1-ACE2 complex, calculated from two parallel experiments in the absence of the peptide, were (1.28 ± 0.01)10^−8^ M and (3.05 ± 0.01)10^−8^ M, respectively. These values agree with the K_D_ = 2.9 x 10^−8^ M, published by Reaction Biology [20], and confirm the reliability of the assay procedure.

**Fig 1.**
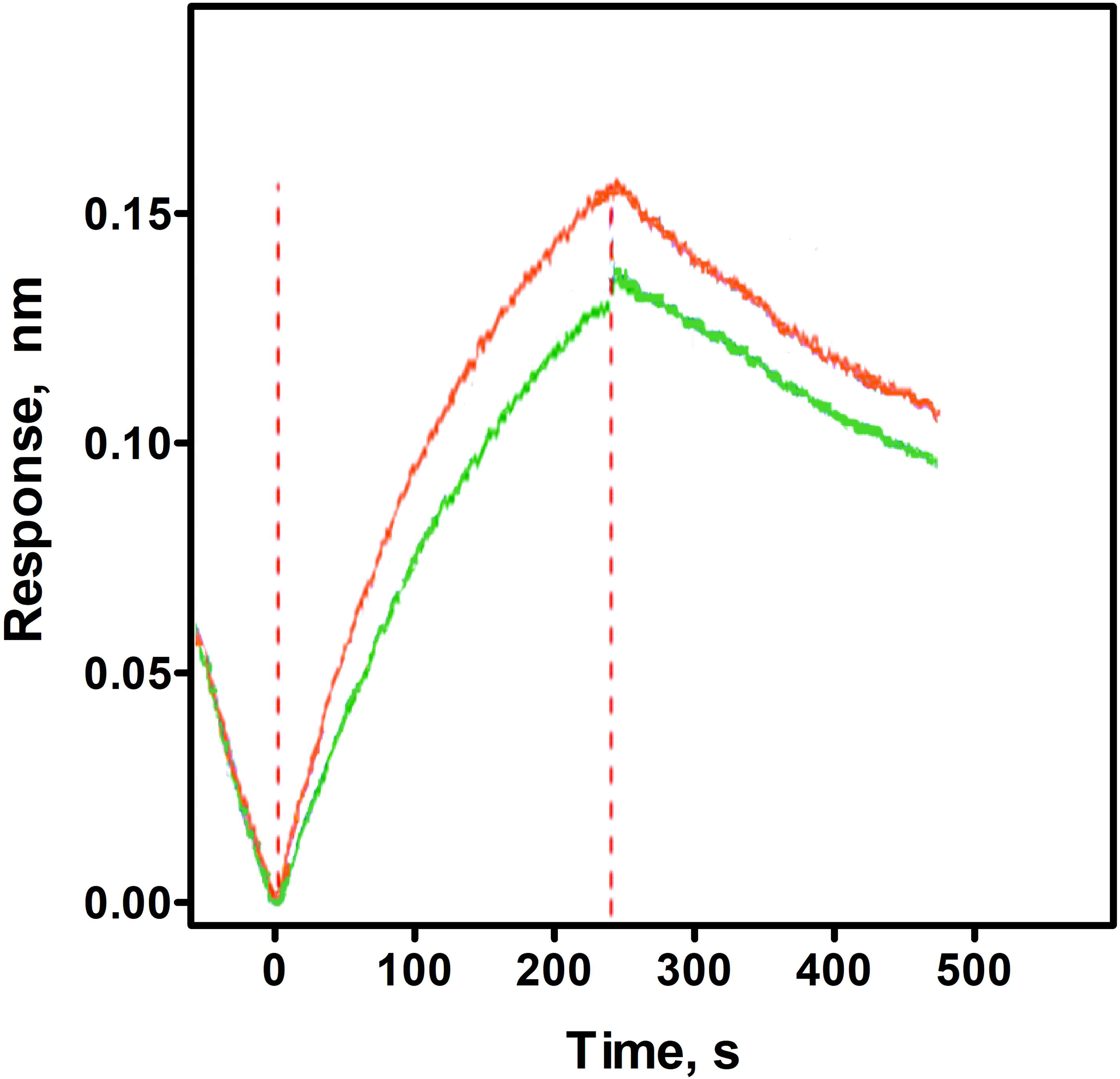
Kinetic curves, characterizing the time course of SARS-CoV-2 spike protein S1 binding to ACE2 protein loaded onto the biosensors of the instrument (ascending curve) and dissociation of this complex (descending curve). Red line: experiment performed using the assay buffer without the peptide. Green line: experiment performed in the presence of 5 μM peptide QAKTFLDKFNHEAEDLFYQ, which had been preincubated with the SARS-CoV-2 spike protein S1 for 15 minutes before the assay.

### Peptide influence on the kinetics of the S1-ACE2 interaction

Fig 1 shows that the time course of interaction of the spike protein S1 with ACE2 (red line) is affected by the addition of 5 μM peptide (green line). For a more detailed analysis of the effect of the peptide on the S1-ACE2 interaction, two series of kinetic experiments were performed, in which varying amounts of the peptide were added to the kinetic assay. One series of experiments simultaneously added the S1 protein and the peptide to the sensor-immobilized ACE2 to initiate the complex formation. In the second series of experiments, preincubation of the S1 protein with the peptide was performed for 15 minutes before initiation of the complex formation. From these data, the k_on_ values were calculated, and the results of this analysis are summarized in Fig 2.

**Fig 2.**
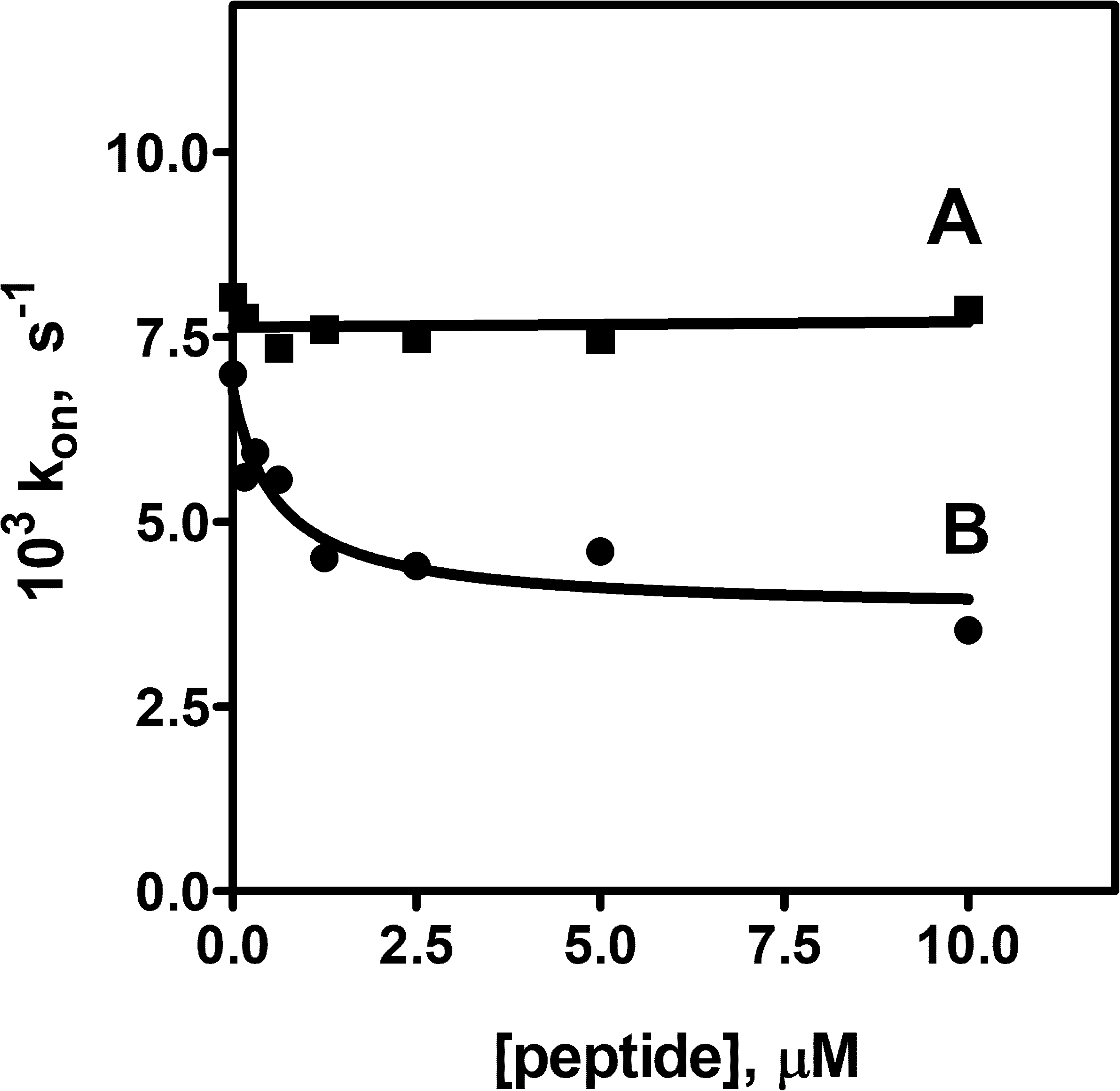
The influence of the peptide QAKTFLDKFNHEAEDLFYQ on the rate constant (k_on_) of S1 binding to ACE2, where ACE2 is immobilized on the biosensor. A. Spike protein S1 and peptide were simultaneously added to the assay buffer before the binding assay was initiated (squares). B. Spike protein S1 was preincubated with the peptide for 15 min in the assay buffer before the binding assay was initiated (circles).

Fig 2 shows that the peptide QAKTFLDKFNHEAEDLFYQ inhibited the rate of S1-ACE2 complex formation, decreasing the rate constant almost two-fold when the spike S1 protein had been preincubated with the peptide. In contrast, no inhibitory effect was observed when the spike S1 protein and peptide were added simultaneously to the assay buffer. These results suggest that the peptide interaction with the spike S1 protein is a slow process, and preincubation is necessary to load the spike S1 protein with the peptide.

Secondly, Fig. 2 shows that the rate constant (k_on_) decreases in the presence of the peptide in a dose-dependent manner, and the half-maximal inhibitory effect was reached at 0.7±0.4 μM. Certainly, this value has some physical meaning if the peptide-spike S1 protein interaction can be described as an equilibrium process.

Finally, the formation of the S1-ACE2 complex was not completely inhibited by the peptide, since the k_on_ values leveled off, even when an excess of the peptide was added. This phenomenon cannot be unambiguously explained with the existing data. However, it appears likely that the incomplete inhibition could be connected to the slow rate of the peptide interaction with its binding site on the S1 protein.

### Dissociation kinetics of the S1-ACE2 complex

Dissociation of the ACE2 bound S1 protein was initiated by transferring the biosensor into fresh assay buffer that did not contain peptide or S1 protein. Therefore, if a binary complex is formed between the ACE2 and S1 proteins, similar k_off_ values, calculated from the descending part of the kinetic curves (see Fig 1), should describe the dissociation process in experiments, performed at different peptide concentration. However, this was not true of this study, as illustrated in Fig 3.

**Fig 3.**
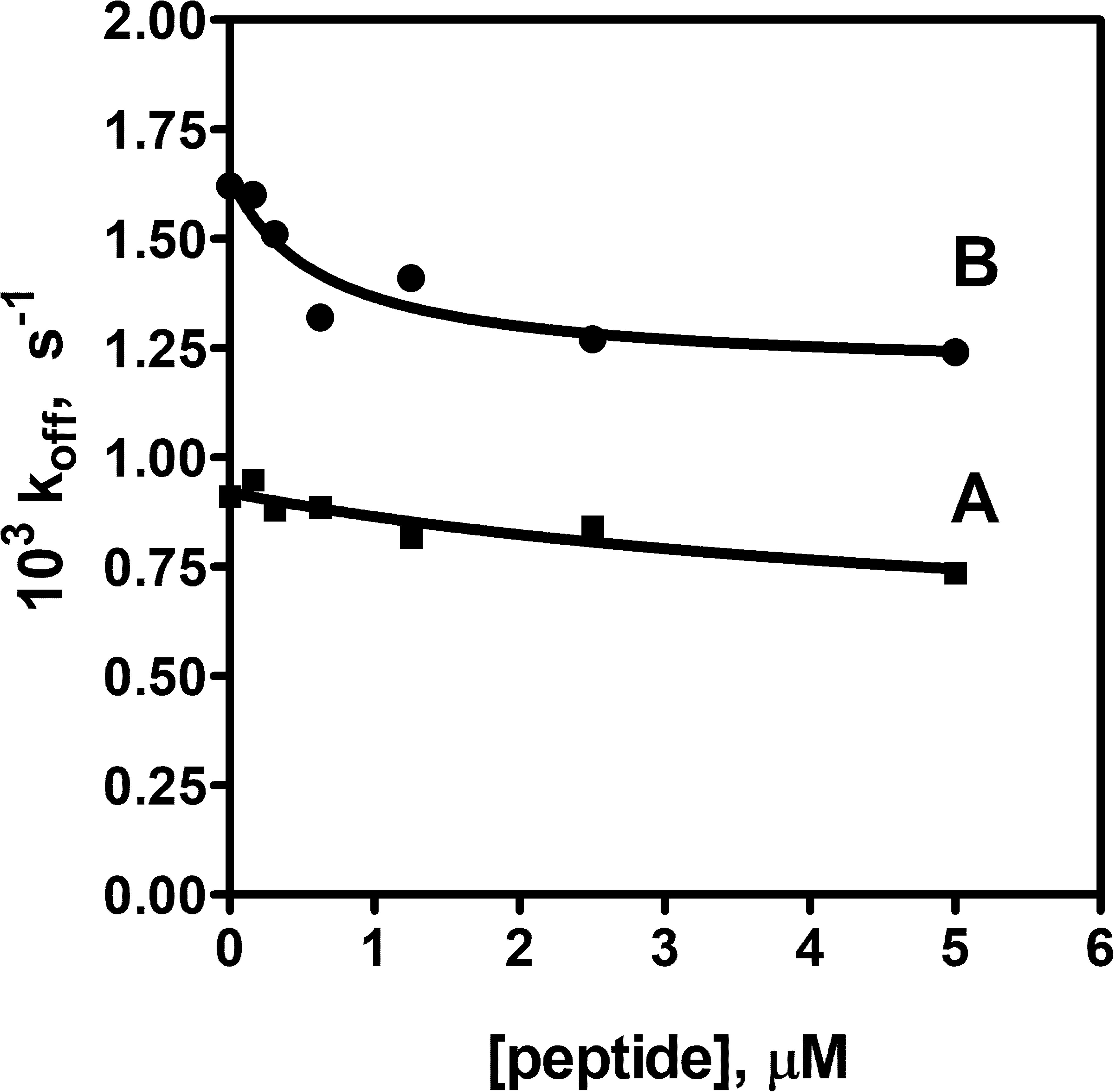
Dissociation of the S1-ACE2 complex, captured by the biosensor in the binding assay. The S1-ACE2 protein complex was formed in the presence of different peptide concentrations in experiments where the S1 protein had been preincubated with the peptide for 15 minutes (series A, squares), or the S1 protein had not been preincubated with the peptide (series B, circles). To initiate the dissociation process, the biosensor was transferred into fresh buffer that did not contain peptide and S1 protein.

It is important to emphasize that dissociation of the S1-ACE2 protein complex was initiated by transfer of the biosensor into buffer that did not contain peptide as well as S1. However, it can be seen in Fig 3 that the k_off_ value depends on the peptide concentration, which was used in the complex formation reaction, demonstrating “memory” relating to the peptide presence in the latter process. These results raise the following questions about the S1-ACE2 complex formation and its structure.

First, the occurrence of such “memory” indicates that different complexes could be formed in the presence of different peptide concentrations, or more likely, the peptide remains in some fraction of this complex, affecting its stability when compared with the binary complex.

Second, half of the peptide’s effect on k_off_ is observed at a peptide concentration of 0.6±0.4 μM, which is in agreement with the effect observed with k_on_ (Fig 2). This may indicate that both effects are caused by the same phenomenon, likely by the formation of the S1-peptide complex. From a chemistry point of view, this is possible if there are several binding sites for this peptide on the S1 protein, as one site must be occupied by the α1 domain of ACE2 in the process of complex formation.

Third, Fig 3 illustrates that preincubation of the peptide with the S1 protein before complex formation with ACE2 destabilized the S1-ACE2 complex since all k_off_ values for series B were higher than the equivalent values in series A. Importantly, this effect did not depend on the peptide presence, as observed when comparing the data points at zero peptide concentration. However, no systematic differences were observed in the binding experiments, performed with both preincubated and non-preincubated S1 protein samples at zero peptide concentration (Fig 2). Thus, the different stabilities of the S1-ACE2 complexes, with or without preincubation of the peptide with S1 protein, seem to be specific for the off-rate reaction, however, the reasons for this phenomenon remain unclear.

Last, although the rate constants k_on_, k_ass_, and k_off_ depend on peptide concentration (Figs 2 and 3), there is practically no influence of the peptide on the K_d_ value, calculated as the ratio of the rate constants k_off_ / k_ass_. Thus, these data demonstrate that even effective peptide interaction with the S1 protein may not shift the equilibrium of the S1 protein binding to ACE2. However, the peptide has a significant effect on S1-ACE2 complex formation and dissociation kinetics. This is an important conclusion to be considered in antiviral therapeutics design, as the simple inhibition mechanism of the virus-receptor binding process by peptides suggested in many papers seems to be oversimplified.

### Alternative peptide binding sites on the S1 protein

The hypothesis that additional (allosteric) binding sites exist on the S1-ACE2 complex, which may bind additional peptide molecules that cause the “memory” effect in the off-rate experiments, was investigated computationally by mapping the putative docking landscape outside the known ACE2 binding site on the S1 protein. These calculations revealed that there are allosteric binding possibilities for the peptide QAKTFLDKFNHEAEDLFYQ (Fig 4).

**Fig 4.**
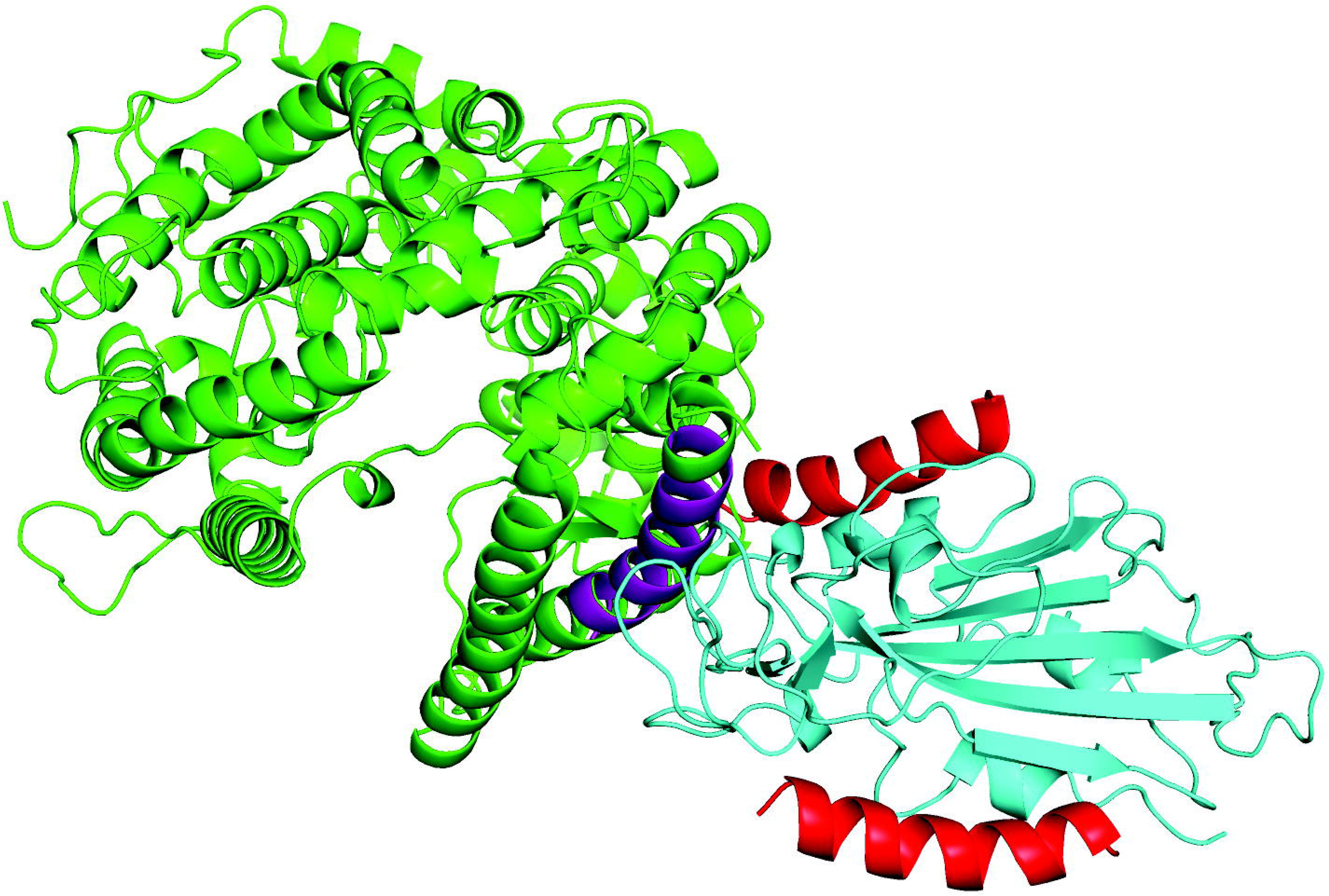
Cartoon representation of the structure of the complex formed between the SARS-Cov-2 spike protein S1 and ACE2. The S1 protein is shown in blue and the ACE2 molecule is shown in green. Two allosterically bound peptide QAKTFLDKFNHEAEDLFYQ molecules are shown in red. The sequence (amino acids 24-42) of the α1 domain of ACE2 which interacts with the S1 protein is shown in violet.

The docking energies of peptide QAKTFLDKFNHEAEDLFYQ at the two allosteric sites were E_dock_=-10.7 kcal/mol (upper location) and E_dock_=−10.5 kcal/mol (lower location), while the docking energy of the same peptide in the recognized RBD of S1 was −11.6 kcal/mol (violet). As these additional sites do not overlap with the RBD, peptide binding to these sites does not necessarily compete with S1 binding to ACE2, however, there may be an allosteric effect. Importantly, the allosteric sites are also available in the free S1 protein, which explains the experimental observations and reasons for the “memory” effect in the off-rate kinetics (where the influence of the peptide used in on-rate experiments is revealed in a dose-dependent manner).

Interestingly, sequential binding of several ACE2 molecules with the spike protein has been discussed in an extensive cryoEM study, where a 1:3 stoichiometry of the spike protein-ACE2 complex was observed [21]. However, if the formation of these multimeric complexes assumes significant conformational changes of participating proteins, binding of additional peptide fragments to the S1 protein appears to be achieved without significant structural changes. Therefore, the physiological meaning for the additional binding sites should be analyzed separately. Furthermore, the presence of several binding sites for the ACE2 α1 domain peptide may open new perspectives for the development of therapeutic agents against SARS-CoV-2 infection.

## Conclusions

The binding kinetics of the spike protein S1 to the SARS-CoV-2 virus receptor ACE2 is affected by the presence of the peptide QAKTFLDKFNHEAEDLFYQ, which corresponds to the sequence 24-42 of the α1 domain of the ACE2 receptor protein. However, as the off-rate of the complex also depends on the concentration of this peptide, which is added to the reaction mixture during the complex formation process, the inhibitory effect of the peptide cannot be clearly observed in the overall binding equilibrium, characterized by the dissociation constant K_d_. The results suggest that the S1 protein has more than one binding site for the ACE2 α1 domain peptide. Our molecular docking calculations confirmed this theory, revealing two other sites, located remotely from the main RBD of the S1 protein. These findings may be important for the development of new peptide-based antiviral therapeutics.

## Acknowledgements

Computational analysis was performed in the High-Performance Computing Center, and the peptide was synthesized in the Core Laboratory of Peptide Chemistry of the University of Tartu. This work was financially supported by QanikDX OÜ, Estonia, registration number 4523084, grant LLTLT20014.

