## Supplementary Figures and Tables for "ACE2 peptide fragment interacts with several sites on the SARS-CoV-2 spike protein S1"

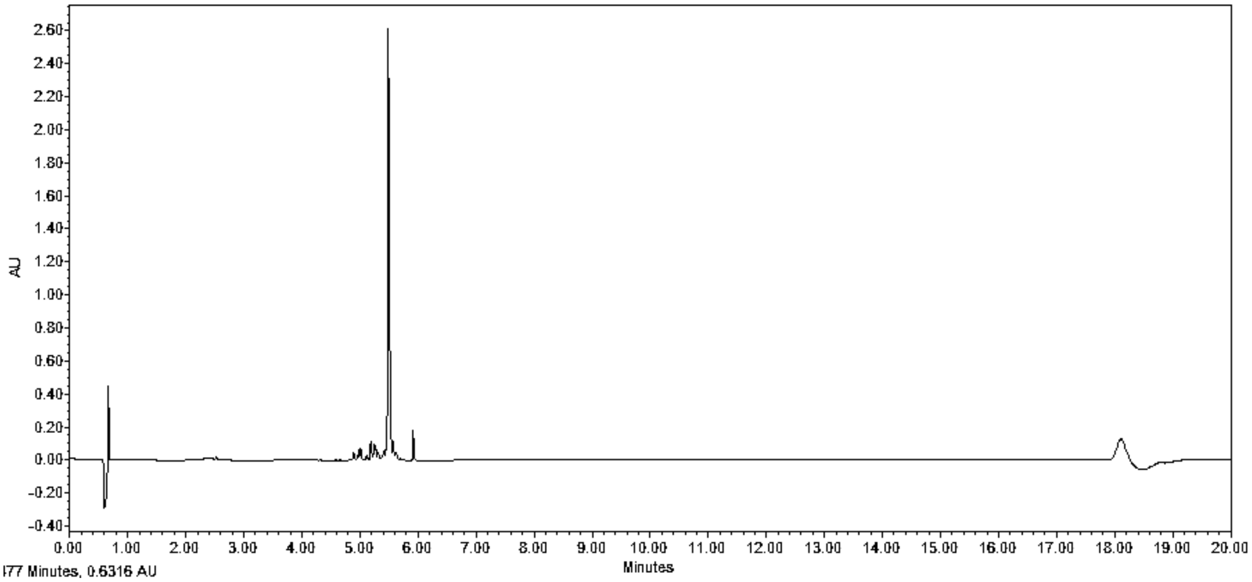


**Fig S1**

**Analysis of peptide QAKTFLDKFNHEAEDLFYQ on a Waters Acquity Ultra-Performance Liquid Chromatography (UPLC) with an acetonitrile/water gradient.** The purity of the peptide was validated at 98%

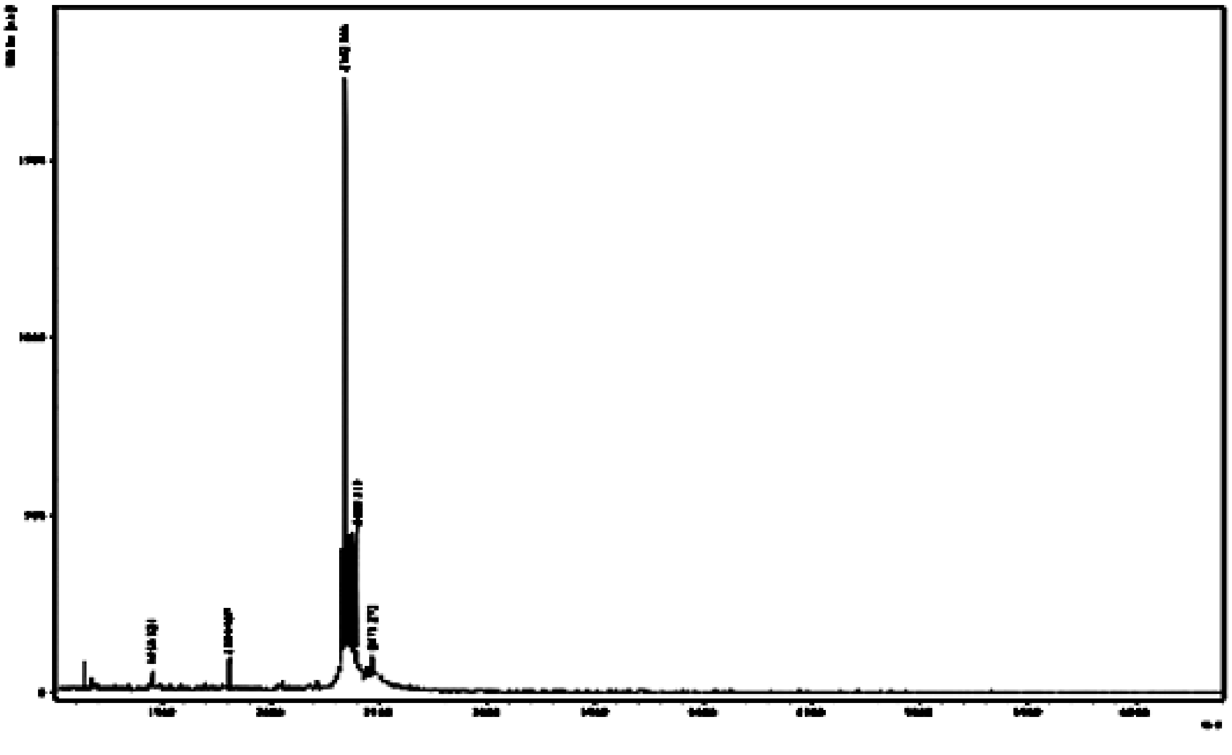

**Fig S2**

**The molecular weight of the peptide QAKTFLDKFNHEAEDLFYQ 2342 kD was determined on a Maldi TOF mass spectrometer Brucker Microflex LT/ SH, USA.**

**Table S1**

**Experiment A**

His-tagged ACE2 was immobilized onto Octet RED96e biosensors (ForteBio, CA, USA) and the binding of S1 protein was measured in the presence or absence of peptide QAKTFLDKFNHEAEDLFYQ. Experiments were performed at 25 °C in 20 mM Tris-HCl pH 7.0 and 150 mM NaCl. Biosensors (HIS1K, lot 6110102) were loaded with His-tagged ACE2, before S1 protein or S1 protein and peptide were added to start the complex formation process.

| Loading Sample ID | Response | $K_D$ (M) | $K_D$ Error | $k_a$ (1/Ms) | $k_a$ Error | $k_{off}$ (1/s) | $k_{off}$ Error |
|---|---|---|---|---|---|---|---|
| P10 uM peptide | 0,1608 | 1,14E-08 | 7,62E-11 | 7,06E+04 | 2,65E+02 | 8,06E-04 | 4,45E-06 |
| P 5 uM peptide | 0,1543 | 1,09E-08 | 1,04E-10 | 6,72E+04 | 3,46E+02 | 7,35E-04 | 5,88E-06 |
| P 2.5 uM peptide | 0,1585 | 1,27E-08 | 8,22E-11 | 6,64E+04 | 2,56E+02 | 8,41E-04 | 4,39E-06 |
| P 1.25 uM peptide | 0,1474 | 1,31E-08 | 1,10E-10 | 6,24E+04 | 3,15E+02 | 8,17E-04 | 5,48E-06 |
| P 0.625 uM peptide | 0,1477 | 1,37E-08 | 7,06E-11 | 6,46E+04 | 2,07E+02 | 8,86E-04 | 3,57E-06 |
| P 0.31 uM peptide | 0,165 | 1,17E-08 | 8,42E-11 | 7,53E+04 | 3,13E+02 | 8,79E-04 | 5,18E-06 |
| P 0.16 uM peptide | 0,1522 | 1,39E-08 | 7,18E-11 | 6,81E+04 | 2,22E+02 | 9,48E-04 | 3,79E-06 |
| Buffer, no peptide | 0,1673 | 1,28E-08 | 1,02E-10 | 7,13E+04 | 3,42E+02 | 9,10E-04 | 5,77E-06 |

| Loading Sample ID | $R_{max}$ | $R_{max}$ Error | $k_{on}$(1/s) | $k_{on}$ Error | $R_{eq}$ | $R_{eq}/R_{max}$(%) | Full X^2 | Full R^2 |
|---|---|---|---|---|---|---|---|---|
| P10 uM peptide | 0,2151 | 0,0005 | 7,87E-03 | 3,09E-05 | 0,1931 | 89,8 | 0,0091 | 0,9974 |
| P 5 uM peptide | 0,2112 | 0,0007 | 7,46E-03 | 4,05E-05 | 0,1904 | 90,1 | 0,015 | 0,9956 |
| P 2.5 uM peptide | 0,2195 | 0,0005 | 7,48E-03 | 3,00E-05 | 0,1949 | 88,8 | 0,0085 | 0,9976 |
| P 1.25 uM peptide | 0,2113 | 0,0007 | 7,55E-03 | 3,70E-05 | 0,1868 | 88,4 | 0,0117 | 0,9964 |
| P 0.625 uM peptide | 0,2073 | 0,0004 | 7,35E-03 | 2,42E-05 | 0,1823 | 87,9 | 0,0048 | 0,9985 |
| P 0.31 uM peptide | 0,2159 | 0,0005 | 7,41E-03 | 3,65E-05 | 0,1933 | 89,5 | 0,0127 | 0,9964 |
| P 0.16 uM peptide | 0,2091 | 0,0004 | 7,76E-03 | 2,60E-05 | 0,1835 | 87,8 | 0,0057 | 0,9982 |
| Buffer, no peptide | 0,2266 | 0,0006 | 8,04E-03 | 4,00E-05 | 0,2009 | 88,7 | 0,0164 | 0,9957 |

**Table S2**

**Experiment B**

His-tagged ACE2 was immobilized onto Octet RED96e biosensors (ForteBio, CA, USA)
and the binding of S1 protein was measured in the presence or absence of peptide QAKTFLDKFNHEAEDLFYQ. Experiments were performed at 25 °C in 20 mM Tris-HCl pH 7.0 and 150 mM NaCl. Biosensors (HIS1K, lot 6110102) were loaded with His-tagged ACE2, before S1 protein or S1 protein and peptide were added to start the complex formation process. In this series of experiments the peptide was preincubated with S1 for 15 min at 25 °C before the binding assay was started.

| Loading Sample ID | Response | $K_D$ (M) | $K_D$ Error | $k_a$ (1/Ms) | $k_a$ Error | $k_{off}$ (1/s) | $k_{off}$ Error |
|---|---|---|---|---|---|---|---|
| S1 & 10 uM peptide | 0,1388 | 4,15E-09 | 2,29E-10 | 3,39E+04 | 4,11E+02 | 1,41E-04 | 7,55E-06 |
| S1 & 5 uM peptide | 0,1385 | 3,59E-08 | 3,54E-10 | 3,45E+04 | 2,99E+02 | 1,24E-03 | 5,86E-06 |
| S1 & 2,5 uM peptide | 0,1406 | 4,06E-08 | 3,56E-10 | 3,13E+04 | 2,46E+02 | 1,27E-03 | 4,90E-06 |
| S1 & 1,25 uM peptide | 0,1308 | 4,53E-08 | 4,13E-10 | 3,11E+04 | 2,59E+02 | 1,41E-03 | 5,20E-06 |
| S1 & 0,625 uM peptide | 0,1377 | 3,10E-08 | 2,25E-10 | 4,25E+04 | 2,62E+02 | 1,32E-03 | 5,04E-06 |
| S1 & 0,31 uM peptide | 0,1484 | 3,42E-08 | 1,86E-10 | 4,43E+04 | 2,10E+02 | 1,51E-03 | 4,06E-06 |
| S1 & 0,15 uM peptide | 0,1375 | 4,00E-08 | 1,63E-10 | 4,00E+04 | 1,46E+02 | 1,60E-03 | 2,88E-06 |
| Buffer, no peptide | 0,1564 | 3,05E-08 | 1,33E-10 | 5,29E+04 | 1,96E+02 | 1,62E-03 | 3,69E-06 |

| Loading Sample ID | $R_{max}$ | $R_{max}$ Error | $k_{on}$(1/s) | $k_{on}$ Error | $R_{eq}$ | $R_{eq}/R_{max}$(%) | Full X^2 | Full R^2 |
|---|---|---|---|---|---|---|---|---|
| S1 & 10 uM peptide | 0,2711 | 0,0025 | 3,53E-03 | 4,86E-05 | 0,2603 | 96 | 0,9946 | 0,9946 |
| S1 & 5 uM peptide | 0,2914 | 0,0019 | 4,69E-03 | 3,57E-05 | 0,2145 | 73,6 | 0,9965 | 0,9965 |
| S1 & 2,5 uM peptide | 0,3169 | 0,002 | 4,40E-03 | 2,95E-05 | 0,2254 | 71,1 | 0,9977 | 0,9977 |
| S1 & 1,25 uM peptide | 0,3012 | 0,002 | 4,51E-03 | 3,11E-05 | 0,2074 | 68,8 | 0,9974 | 0,9974 |
| S1 & 0,625 uM peptide | 0,253 | 0,0011 | 5,57E-03 | 3,12E-05 | 0,1932 | 76,4 | 0,9973 | 0,9973 |
| S1 & 0,31 uM peptide | 0,2709 | 0,0009 | 5,94E-03 | 2,51E-05 | 0,2019 | 74,5 | 0,9983 | 0,9983 |
| S1 & 0,15 uM peptide | 0,2659 | 0,0007 | 5,60E-03 | 1,75E-05 | 0,19 | 71,5 | 0,9992 | 0,9992 |
| Buffer, no peptide | 0,2536 | 0,0006 | 7,30E-03 | 2,32E-05 | 0,1943 | 76,6 | 0,9985 | 0,9985 |
